# Geometry of Ranked Nearest Neighbour Interchange Space of Phylogenetic Trees

**DOI:** 10.1101/2019.12.19.883603

**Authors:** Lena Collienne, Kieran Elmes, Mareike Fischer, David Bryant, Alex Gavryushkin

## Abstract

In this paper we study the graph of ranked phylogenetic trees where the adjacency relation is given by a local rearrangement of the tree structure. Our work is motivated by tree inference algorithms, such as maximum likelihood and Markov Chain Monte Carlo methods, where the geometry of the search space plays a central role for efficiency and practicality of optimisation and sampling. We hence focus on understanding the geometry of the space (graph) of ranked trees, the so-called ranked nearest neighbour interchange (RNNI) graph. We find the radius and diameter of the space exactly, improving the best previously known estimates. Since the RNNI graph is a generalisation of the classical nearest neighbour interchange (NNI) graph to ranked phylogenetic trees, we compare geometric and algorithmic properties of the two graphs. Surprisingly, we discover that both geometric and algorithmic properties of RNNI and NNI are quite different. For example, we establish convexity of certain natural subspaces in RNNI which are not convex is NNI. Our results suggest that the complexity of computing distances in the two graphs is different.

## 1. Introduction

The nearest neighbour interchange (NNI) graph is defined on the set of phylogenetic trees with adjacency relation given by the interchange operation of two sister clades (subtrees). It has been known in mathematical biology literature for nearly 50 years (Robinson 1971; Moore, Goodman and Barnabas 1973). Its metric geometry has been extensively studied (Dasgupta et al. 2000; Li, Tromp and Zhang 1996; Gordon, Ford and St John 2013; Jong, McLeod and Steel 2016). An important property of the NNI graph is that computing distances is NP-hard (Dasgupta et al. 2000), a fact that goes partway towards explaining why tree search and sampling algorithms pose a significant challenge even for moderately sized trees (Whidden and Matsen 2016).

Recent advances in computational phylogenetics have introduced various classes of molecular clock models (Yoder and Yang 2000; Drummond, Ho et al. 2006; Drummond and Suchard 2010) and made computational inference of phylogenetic time-trees possible (Ronquist and Huelsenbeck 2003; Bouckaert et al. 2018; Hadfield et al. 2018). However, the mathematical challenges resulting from this seemingly minor change in parametrisation (genomic distance versus time distance) of trees have only recently been brought to attention (Gavryushkin and Drummond 2016). These differences motivated Gavryushkin, Whidden and Matsen (2018) to propose an extension of the NNI graph to the class of discrete time-trees. The simplest such extension introduces the RNNI graph on the set of ranked phylogenetic trees. Two rooted trees are considered adjacent if they differ by a single neighbour interchange or order change respecting that ranking (detailed description below). As a metric space, the RNNI graph inherits the geometric and algorithmic challenges that the NNI space has been traditionally facing. Surprisingly, most of them cannot be settled by directly translating results or applying techniques developed for NNI (Gavryushkin, Whidden and Matsen 2018).

In this paper, we consider the RNNI space on ranked phylogenetic trees which have all taxa being of equal rank zero. An example of such a tree is depicted in Figure 1. In the terminology of (Gavryushkin, Whidden and Matsen 2018), the space considered in this paper is the space of ranked ultrametric phylogenetic trees (RNNIu in their notation). We postpone formal definitions until later in this section.

**Figure 1.**
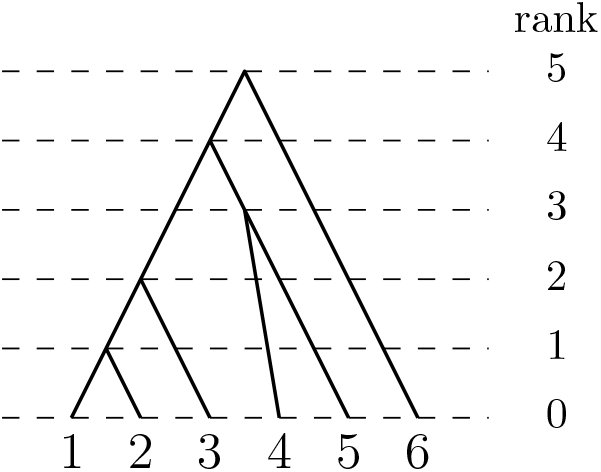
A ranked tree with 6 taxa.

We investigate the geometry and algorithmic complexity of the RNNI metric space. Specifically, in this paper we establish the exact radius and diameter of the space (Section 3.2). We also show that the subset of caterpillar trees is convex in the RNNI space (Section 3.1), thus settling one of the open problems proposed in (Gavryushkin, Whidden and Matsen 2018). To establish these geometric properties we use algorithms that will be introduced in Section 2. We will in particular provide an approximate algorithm that computes exact distances for small trees. The question of whether there exists a polynomial algorithm for computing the RNNI distance remains an open problem.

In the remainder of this section we formally introduce the terminology used in this paper.

A *ranked phylogenetic tree* is a pair consisting of a rooted binary phylogenetic tree *T* on the set *X* = {1, …, *n*} of *taxa* for 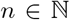, and a (total) rank function that maps all leaves of *T* to 0, all internal nodes of *T* onto elements of the set {1, …, *n* −1}, and respects the partial order on the nodes given by the tree. The latter means that if *u* and *v* are two internal nodes of *T* such that there exists a path from a taxon *x ∈ X* to the root which first passes through *u* and then through *v* then rank(*u*) *<* rank(*v*). This definition also implies that the rank of every internal node is distinct and every *i* with 1 ≤ *i* ≤ *n* − 1 equals the rank of some node. Ranked trees (*T*_1_, rank_1_) and (*T*_2_, rank_2_) are considered equal if *T*_1_ = *T*_2_ and rank_1_ = rank_2_. Since all trees in this paper are ranked trees, we will abuse the notation and drop the rank function from the notation. We will also simply say trees to mean ranked phylogenetic trees. For a tree *T*, we will use rank_*T*_ to refer to its ranking. A ranked tree *T* without its leaf labels will be called *tree topology*.

Our definition of a ranked tree implies a natural notion of edge length – we call the difference in rank |rank_*T*_ (*v*)−rank_*T*_ (*u*) the length of an edge (*u, v*). We assume that edges of trees are undirected, so (*u, v*) = (*v, u*).

Now we are ready to introduce the tree space which is the subject of study in this paper, the RNNI graph.

The vertex set of the RNNI graph is the set of all ranked trees on *n* taxa. We introduce two types of operation (RNNI *moves*) on trees (see Figure 2) and say that two trees are adjacent in the RNNI graph if one can be obtained from the other by an operation of either type. The first type of operation is called a *rank move* and defined by swapping the ranks of two internal nodes that are not adjacent in the tree but are consecutive in the ranking. Formally, if *u* and *v* are nodes of a tree *T* such that |rank_*T*_(*u*) rank_*T*_(*v*)| = 1 and (*u, v*) is not an edge in *T* then the tree *R* obtained from *T* by only changing rank_*R*_(*u*) = rank_*T*_ (*v*) and rank_*R*_(*v*) = rank_*T*_ (*u*) is said to be obtained by a rank move. The second type of operation is called an NNI *move* and defined in the usual way, that is, two trees *T* and *R* are said to be connected by an NNI move if there exist internal edges *e* in *T* and *f* in *R* both of length one such that the trees obtained by shrinking *e* and *f* to internal nodes coincide.

**Figure 2.**
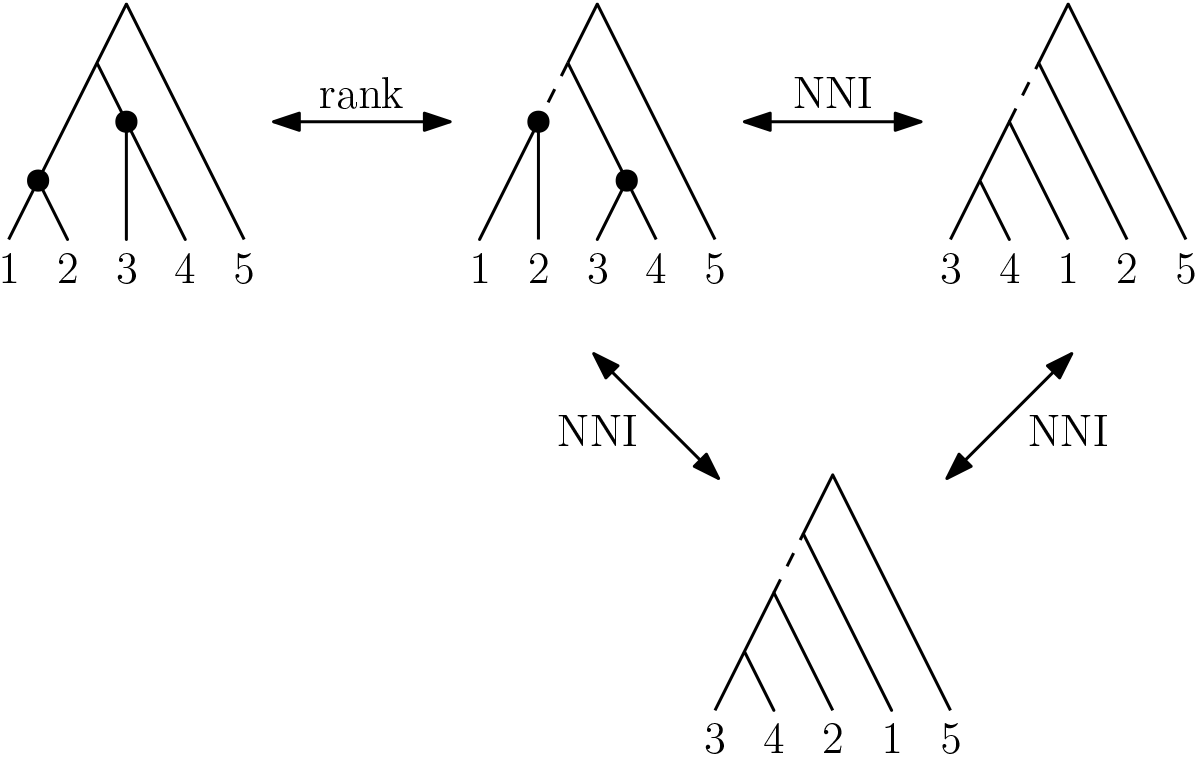
Two types of operation that define edges of the RNNI graph. The rank move swaps the rank of the two highlighted nodes and the NNI moves are performed on the dashed edges.

We use the notation *d*(*T, R*) throughout the paper to denote the length of a shortest RNNI path between tree *T* and *R*. An important geometric property analysed in this paper is whether trees from a certain class are connected but shortest paths that do not leave the class. We say that a subset *S* of trees is *convex* if every pair of trees from *S* is connected by a shortest path that completely stays within *S*.

The rest of this paper is organised as follows. In Section 2 we introduce three algorithms for exploring the RNNI graph. We use these algorithms in Section 3 to establish several geometrical properties of this graph, such as its diameter and radius. Furthermore, we will establish in Section 3.1 that a certain subspace of RNNI is convex and design a polynomial-time algorithm for computing RNNI-distances in that subspace. This result suggests that the complexity of computing distances in RNNI is different from that in the classical NNI space. We will discuss this further in Section 3.3, where we disprove a conjecture of Gavryushkin, Whidden and Matsen (2018) and suggest weaker versions as plausible alternatives. In Section 3.4 we establish a connection between the RNNI graph and a classical algebraic structure, the partition lattice. We will finish the paper with a discussion and open problems.

## 2. Algorithms

In this section we introduce three algorithms, FindPath (Section 2.1), Caterpillar Sort (Section 2.2), and MDTree (Section 2.3). FindPath computes an RNNI path between trees. Caterpillar Sort computes a shortest RNNI path between caterpillar trees (also known as ladder trees, see later in this section for formal definitions). MDTree attempts to maximise the dissimilarity to find a remote tree from an arbitrary RNNI tree. While these algorithms are interesting on their own (e.g. in simulation studies), they will also form an important ingredient of our study of the geometry of the RNNI space later in this paper. For example, MDTree will be an important tool for finding the radius of the RNNI graph, as the algorithm computes a tree with maximum distance if the input tree is a caterpillar tree. We will prove this fact in Section 3.2.

The algorithm FindPath provides an upper bound the RNNI distance. We will study the accuracy of this approximation by also computing the exact RNNI distance for small trees. Specifically, we will use the algorithm for computing the complete RNNI graph that is given in (Gavryushkin, Whidden and Matsen 2018, Section 3.3). After pre-computing the graph, we use Seidel’s algorithm (Seidel 1995) for computing distances. However, the large number of vertices in RNNI is an obstacle when it comes to computing distances for larger trees. Since there are 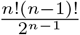 vertices in the RNNI graph (Gavryushkin, Whidden and Matsen 2018), we compute the RNNI graph on up to seven taxa for our analyses. This computational result will also be used later in the paper to support Conjecture 10, a modification of Conjecture 9 in (Gavryushkin, Whidden and Matsen 2018), which we prove to be false in this paper.

Note that computing the RNNI space on trees with seven taxa is a challenging goal. For example, Whidden and Matsen (2016) were able to compute the much smaller SPR space on up to seven taxa and used this to compute curvature values for all pairs of trees within that space. We have implemented all our algorithms in a combination of Python and C, and the source code is available at (Collienne, Elmes and Gavryushkin 2019).

### 2.1 FindPath

In this section we present FindPath, an efficient heuristic for computing short paths between trees in RNNI.

Before introducing the algorithm we need the following definitions. Each node *v* of a tree defines a *cluster C*, which is the set of taxa descending from *v*. We then say that *v induces C*. All trees on *n* leaves share trivial clusters, which are those induced by leaves and the root, so we exclude them from consideration and assume cluster means a non-trivial cluster. A tree is unambiguously defined by the list of its clusters sorted according to the rank of the inducing nodes. We will refer to this list of *n* − 2 clusters for a given tree on *n* taxa as a *cluster representation* of the tree (Gavryushkin, Whidden and Matsen 2018; Gavryushkin and Drummond 2016; Semple and Steel 2003).

Let *T* and *R* be arbitrary trees and [*C*_1_, …, *C*_*n*−2_] be the cluster representation of *R* (the “destination” tree). Then *C*_*i*_ is induced by the node of rank *i* for *i* = 1, …, *n* − 2. FindPath computes a path by iteratively extending a sequence *p* of trees starting from *T*. The algorithm terminates when *p* is an RNNI path from *T* to *R*. At step *k* = 1, …, *n* − 2 of FindPath we consider the cluster *C*_*k*_ and extend path *p* as follows. We search for a node in the last tree *T′* of *p* from the previous step that induces the smallest cluster containing all elements of *C*_*k*_. This node is denoted by mrca_*T′*_ (*C*_*k*_) (short for *most recent common ancestor*). We then extend *p* by adding the tree obtained from *T′* by performing an RNNI move that decreases the rank of mrca_*T′*_ (*C*_*k*_). We continue extending *p* in this way by repeatedly performing RNNI moves until the rank of mrca_*T′*_ (*C*_*k*_) = *k*, thus completing step *k*.

#### Algorithm 1 FindPath(*T, R*)

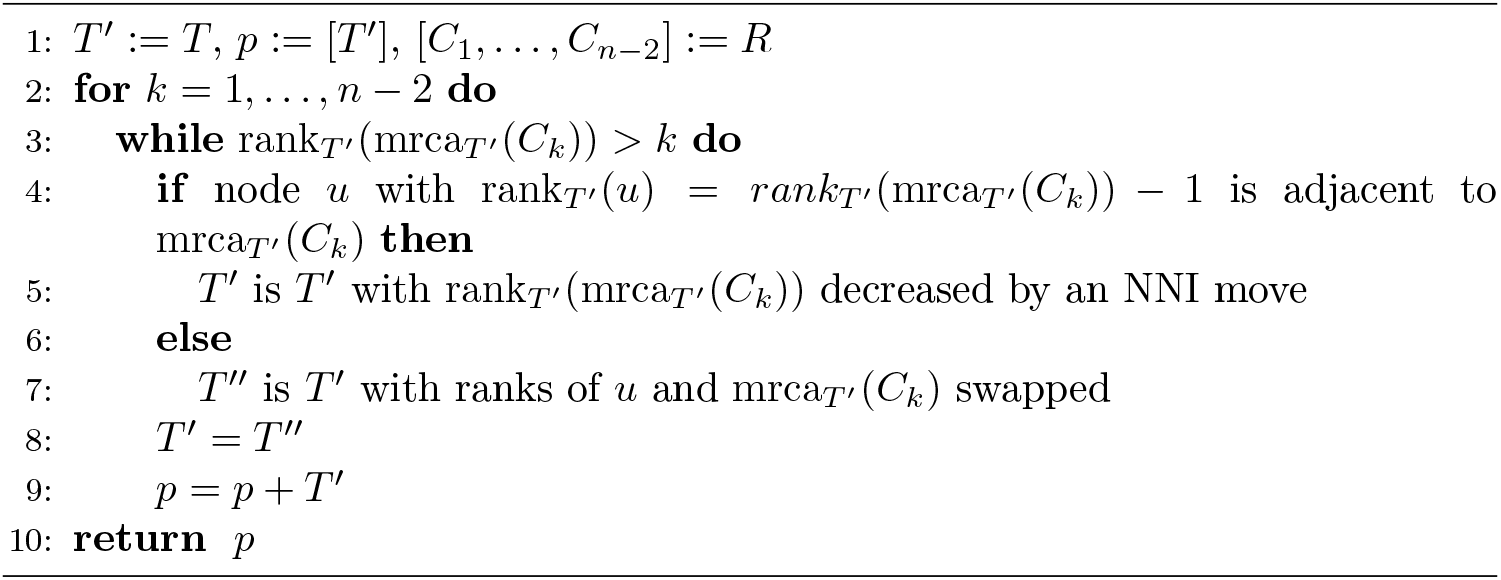

**Proposition 1.** FindPath *is a correct deterministic algorithm*.

*Proof*. It is not hard to see that the algorithm terminates with a path ending in *R*. Indeed, every iteration of the **for** loop (line 2) is completed once rank_*T′*_ (mrca_*T′*_ (*C*_*k*_)) = *k*, that is, the first *k* clusters of *T′* and *R* coincide. Hence after *n* − 2 such iteration the path has to arrive at *R*.

To complete the proof, it suffices to show that the update operation (line 7) is well-defined, that is the RNNI move that decreases the rank of mrca_*T′*_ (*C*_*k*_) is unique.

Case *k* = 1. In this case *C*_*k*_ consists of two taxa {*x, y*}. The node *v* = mrca_*T′*_ (*x, y*) has rank_*T′*_ (*v*) = *r >* 1. Consider the node *w* preceding *v* in *T′*, that is, rank_*T′*_ (*w*) = *r* − 1. If a rank move that swaps *v* and *w* is possible then this is the only move on *T′* that can decrease the rank of mrca(*x, y*). If the rank move is impossible then there is an edge in *T′* connecting *v* and *w*. The only way to decrease the rank of mrca(*x, y*) then is to perform an NNI move on that edge. It is not hard to see that out of two possible NNI moves only one decreases the rank of this mrca.

Case *k >* 1.

1. *C*_*k*_ = *C*_*i*_ ∪ *C*_*j*_ for *i, j < k*. Suppress *C*_*i*_ and *C*_*j*_ in both *T′* and *R* to new taxa *c*_*i*_ and *c*_*j*_ respectively and proceed as in Case *k* = 1.
2. *C*_*k*_ = *C*_*i*_ ∪ {*x*}, where *x* is a taxon not present in *C*_1_, …, *C*_*k*_. Suppress *C*_*i*_ in both *T′* and *R* to a new taxon *c*_*i*_ and proceed as in Case *k* = 1.
3. *C*_*k*_ = {*x, y*} where both *x* and *y* are not in *C*_1_, …, *C*_*k*_. Proceed as in Case *k* = 1.

Clearly, the worst-case complexity of FindPath is quadratic in the number of taxa.

Because the algorithm returns a path between pairs of trees in RNNI, the length of the path approximates the RNNI distance from above. A natural question then is how accurate this approximation is. We have computationally shown that the algorithm FindPath finds the correct distance for trees up to seven taxa (Collienne, Elmes and Gavryushkin 2019) and know of no larger counterexample.

### 2.2 Caterpillar Sort

In this section we introduce an algorithm to compute RNNI paths between *caterpillar trees*, which are trees where each internal node is adjacent to at least one leaf. Caterpillar trees have only one *cherry*, which is a pair of taxa that share their parent. A path between two caterpillar trees that only consists of caterpillar trees is called a *caterpillar path*. We can identify a caterpillar tree *T* = [{*x*_1_, *x*_2_}, {*x*_1_, *x*_2_, *x*_3_}, …, *x*_1_, …, *x*_*n*−1_}] with the list of its taxa [*x*_1_, *x*_2_, …, *x*_*n*_], assuming that *x*_1_ < *x*_2_ (recall that the set of taxa is {1, …, *n*}).

The algorithm Caterpillar Sort (Algorithm 2) is a modification of the classical Bubble Sort algorithm (Knuth 1997). A path *p* from *T* to *R* = [*x*_1_,…, *x*_*n*_] is computed iteratively so that after *k* steps the last *k* taxa of *T* and *R* coincide. Specifically, in step *k* (*k* = *n*, …, 3) the parent of taxon *x*_*k*_ is moved up by NNI moves to the position it has in *R*. Notice that there might be more than just one such move per step, which means that *p* can be extended by more than one tree in each step.

#### Algorithm 2 Caterpillar Sort(*T, R*)

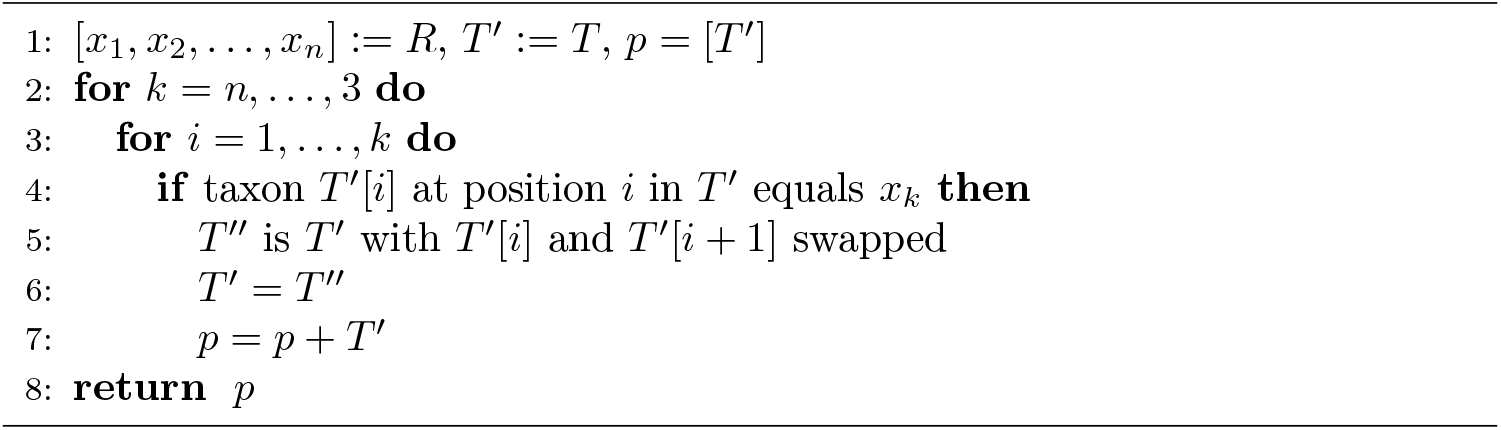

The running time of Caterpillar Sort is quadratic in *n*.

The path *p* returned by Caterpillar Sort is a shortest caterpillar path because every tree modification on *p* reduces the number of inversions of taxa in *T* and *R*, and no modification can reduce more than one insertion. Indeed, no RNNI move on caterpillar trees can reduce the number of taxa inversions by more than one, assuming that inversions with both taxa of a cherry are counted only once. That is, if both pairs of taxa (*x, z*) and (*y, z*) appear in opposite orders in *T* and *R* and *x* an *y* form a cherry then they are counted as one inversion. Notice that because of this the distance between caterpillar trees does not coincide with the Kendall tau distance (Kendall 1948), counting the number of pairs of elements that appear in different order in two permutations, even though the idea is very similar. Hence any caterpillar path, every move along which reduced the number of inversions, is a shortest caterpillar path.

It is not obvious that the path between *T* and *R* returned by Caterpillar Sort has the least possible length among all RNNI paths (not only caterpillar paths), that is, the length of *d*(*T, R*). This fact will be established in Theorem 5.

Our Python implementation of this algorithm can be accessed at (Collienne, Elmes and Gavryushkin 2019).

### 2.3. MDTree

The idea behind this algorithm is to efficiently return a tree as far away from a given tree as possible. As we will see in Section 3.2, MDTree achieves this goal for caterpillar trees.

MDTree (Algorithm 3) works as follows. Given an arbitrary tree *T*, order its taxa in a list *L* = [*l*_1_, …, *l*_*n*_] so that the ranks of their parents are non-decreasing with respect to this order. Note that this list is not uniquely determined by the tree. MDTree constructs an output tree *R* as follows: Initially, *R* only consists of two taxa *l*_1_ and *l*_2_, the most recent common ancestor of which will eventually be of rank *n* − 1. The remaining taxa are iteratively added to *R* reversing the order of *L*. At every iteration, the taxon is attached to a new internal node created on one of the existing branches in *R* so that this new attachment node has rank one. Note that *R* is not uniquely determined due to the non-deterministic choice of the attachment edge.

#### Algorithm 3 MDTree(*T*)

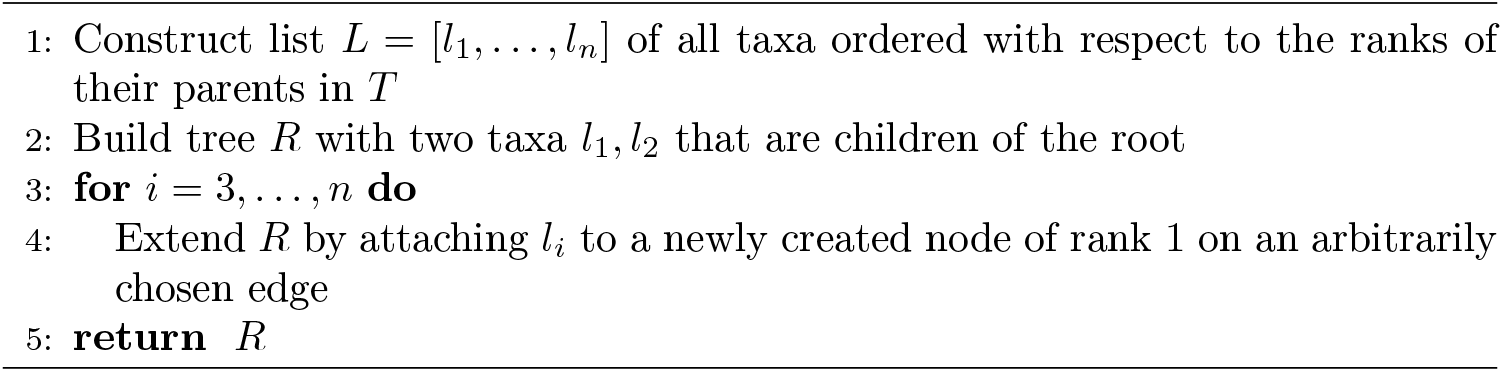

Note that the list computed in Line 1 of MDTree, as well as the attachment edge of the new taxon in Line 4 are not uniquely determined. Therefore, this algorithm is non-deterministic, which is an important observation that we will use for finding the radius of RNNI in Section 3.2.

## 3. Geometry of RNNI

The study of shortest paths in the RNNI graph in this section is motivated by our aim to understand the geometry of RNNI as a metric space. We will employ the algorithms developed in the previous section to aid our analysis of shortest paths. Those algorithms will, for example, enable us to prove that the set of caterpillar trees is convex in RNNI (Theorem 5 in Section 3.1). In Section 3.2 we will investigate the diameter and radius of the RNNI space. Interestingly, the *diameter* of RNNI, that is the maximum distance ∆(RNNI) = max {*d*(*T, R*) | *T, R* ∈ RNNI} between any two trees in the graph, equals its *radius* defined as 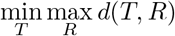. Next, we will consider the so-called Split Property (Gavryushkin, Whidden and Matsen 2018), that is, every shared split of taxa is maintained along every shortest path. Recall that a *split* is a bipartition of the set of taxa obtained by deleting an edge of the tree *T*. Splits are denoted by *A|B*. We will give a counterexample to show that RNNI does not have the Split Property and consider a variant, the Weak Split Property, which says that every shared split of taxa is maintained along a shortest path. We will also consider the so-called Cluster Property as another weak version of the Split Property (see Conjecture 10). In the final part of this geometry section we will discuss the relation between the RNNI graph and the so-called partition lattice, a well-known algebraic structure.

We start with the following auxiliary results.

### Lemma 2. 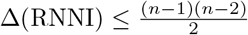 *for n ≥* 3

*Proof.* Since the diameter is bounded from above by the maximum length of a path computed by FindPath, it is enough to find this maximum. Let us assume that *T* and *R* = [*C*_1_, …, *C*_*n*−2_] are trees for which FindPath computes a path of maximum length. It follows that in each step of FindPath the most recent common ancestor of the cluster *C*_*k*_ considered in that step is the root. Thus, there are *n* − 1 − *k* RNNI operations necessary to move *C*_*k*_ to its correct position. Hence, the maximum length of a path computed by FindPath is bounded by 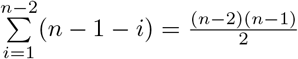. □

The following lemma relates distances between trees on *n* and *n* + 1 taxa and is an important tool for inductive arguments. We use the notion parent_*T*_(*x*) to refer to the node adjacent to taxon *x* in tree *T*. Let *T*|_*n*_ denote the restriction of tree *T* to the set of taxa {1, …, *n*}. In other words, if *T* is a tree on taxa {1, …, *n* + 1} the tree *T*|_*n*_ is obtained by deleting taxon *n* + 1 and suppressing the thereby created node of degree two, updating node rankings accordingly.

### Lemma 3. *Let T and R be two trees on taxa* {1, …, *n* + 1}. *Then d*(*T*|_*n*_,*R*|_*n*_) ≤ *d*(*T, R*) − *δ*, *where δ* = |rank_*T*_ (parent_*T*_ (*n* + 1)) *−* rank_*R*_(parent_*R*_(*n* + 1))|

*Proof*. First observe that the rank of the internal node parent_*T*_ (*n* + 1) can only be changed by performing a rank move that involves parent_*T*_ (*n* + 1) or an NNI move on an edge adjacent to parent_*T*_ (*n* + 1). Second observe that any RNNI move can change the rank of parent_*T*_ (*n* + 1) by at most one. Hence an RNNI move on *T* can decrease the rank of parent_*T*_ (*n* + 1) by at most one.

Let *p* be a shortest path from *T* to *R* and *p*|_*n*_ the path resulting from deleting taxon *n* + 1 from all trees on *p* and removing all identical trees. Then *p*|_*n*_ is a path from *T*|_*n*_to *R*|_*n*_. Recall that *δ* is the difference in ranks between the parents of taxon *n* + 1 in *T* and *R*. Combining this with the observations above, we conclude that |*p*|_*n*_| ≤ |*p*| − *δ*. Since *d*(*T*|_*n*_, *R*|_*n*_) ≤ |*p*|_*n*_| and |*p*| = *d*(*T, R*), the desired inequality follows. □

### 3.1. The set of caterpillar trees

In this section we restrict our attention to the set of caterpillar trees. These trees are of particular interest because they are used to prove that computing NNI distances is an NP-hard problem. As we will see below, shortest paths between caterpillar trees in the RNNI space differ from those in the classical NNI space. We will show later in this section that RNNI distances between caterpillar trees can be computed in polynomial time. Throughout this section we use the list representation of caterpillar trees described in Section 2.2.

Gavryushkin, Whidden and Matsen (2018) showed that computing a shortest path between two caterpillar trees in the NNI graph sometimes requires first building a clade and then moving the clade around the tree (see Figure 3). In the RNNI graph however, the necessity of additional rank moves invalidates this strategy. This is due to the fact that NNI moves in RNNI are only allowed on edges of length one.

**Figure 3.**
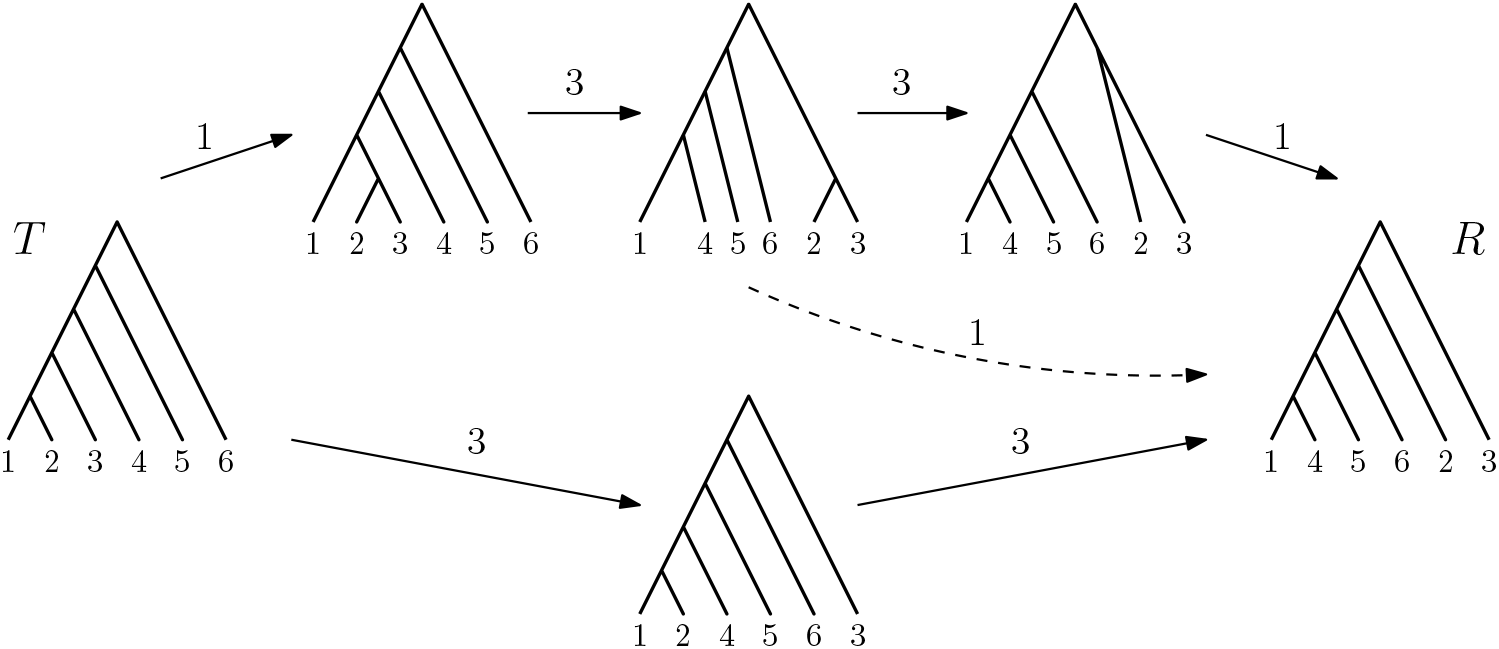
Paths between caterpillar trees *T* and *R*: Solid arrows indicate paths in RNNI, the dashed arrow is a move only possible in NNI. The bottom path is a shortest RNNI path.

These observations motivate the following general result. We will show that every pair of caterpillar trees *T, R* in RNNI are connected by a caterpillar path of length *d*(*T, R*), that is, the set of caterpillar trees is convex in RNNI (Theorem 5). Note that this is not true in the NNI space (Figure 3).

For proving the convexity of the set of caterpillar trees we need the following lemma. Recall that 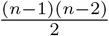 is an upper bound of the diameter of RNNI by Lemma 2.

#### Lemma 4. *Let T and R be two caterpillar trees. Then* 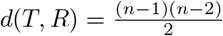 *if and only if* 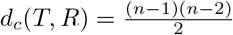, *where d*_*c*_ *is the length of a shortest caterpillar path*

*Proof*. First note that 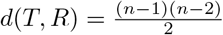 implies 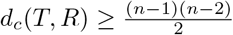. In each step *i* = 0, …, *n* 3 of Caterpillar Sort at most *n* − 2 − *i* pairs of taxa swap positions, as in the worst case the taxon considered in step *k* is moved from the cherry of the tree up to the position *n − k*. It follows that the maximum length of a path computed by Caterpillar Sort is

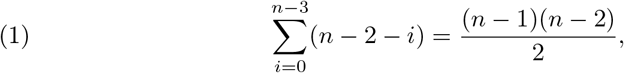

so 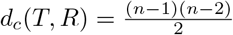

To prove the converse, assume that 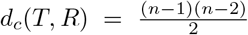. We prove that 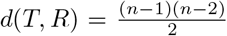 by induction on the number of taxa *n*. The induction basis for *n* = 3 taxa is trivial as the RNNI graph is a triangle on three caterpillar trees in this case.

For the induction step we assume without loss of generality that *T* is the caterpillar tree [1, …, *n* + 1] and *R* is a caterpillar tree such that 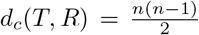. Consider the path *p* = Caterpillar Sort(*T, R*) of this length. Note that in this case taxon *n* + 1 has to be in the cherry of *R*, as otherwise Caterpillar Sort would find a path between *T* and *R* of length strictly less than 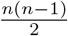, which can be shown by counting the number of moves as in (1) above.

Let us now consider the path *p*|_*n*_ from *T*|_*n*_to *R*|_*n*_, which is obtained by restricting every tree on *p* to taxa 1, …, *n* and removing identical trees. Observe that this path is shorter than *p* by the number of moves that involve taxon *n* + 1. Since this taxon is adjacent to the root in *T* and part of the cherry in *R*, Caterpillar Sort requires *n −* 1 such moves. Hence, 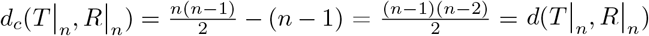, by the induction hypothesis. Lemma 3 implies that *d*(*T*|_*n*_, *R*|_*n*_) ≤ *d*(*T,R*) − |rank_*T*_(parent_*T*_(*n* + 1)) − rank_*T*_(parent_*T*_(*n* + 1))| = *d*(*T, R*) − (*n* − 1). So 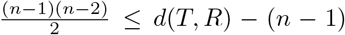, which implies that 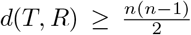. Thus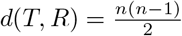. □

#### Theorem 5. *The set of caterpillar trees is convex in* RNNI

*Proof*. Note that it suffices to prove that *d*_*c*_(*T, R*) = *d*(*T, R*) for an arbitrary pair of caterpillar trees *T* and *R*. Lemma 4 implies that if *T* and *R* are at the maximal possible distance, it is 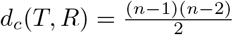. Denote the latter number by *D*.

Assume now that *d*_*c*_(*T, R*) = *D* − 1. If we apply Caterpillar Sort(*T, R*) in this case, there must be a step in the algorithm where the taxon that is moved up is not part of the cherry (of *T′*) at the beginning of this step. Let *x* be the first taxon that has this property and is considered at step *k*. Then there must be a taxon *y* that is immediately preceding *x* in the tree at the beginning of step *k*. Since *k* is the first such step, *y* must be preceding *x* in tree *T* as well. Consider a tree 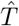 that is obtained from *T* by exchanging *x* and *y*. Note that the caterpillar path from 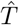 to *R* computed by Caterpillar Sort passes *T* and coincides with the path Caterpillar Sort(*T, R*) from there on. Clearly, 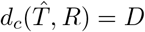.

Assuming *d*_*c*_(*T, R*) < *D*, we can iterate the construction above to find a tree 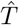 such that 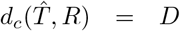 and the caterpillar path from 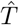 to *R* computed by Caterpillar Sort passes *T* and coincides with the path Caterpillar Sort(*T, R*) from there on. Lemma 4 implies that 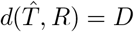.

Assume that *d*(*T, R*) < *d*_*c*_(*T, R*) and let *p* be a path from *T* to *R* of length *d*(*T, R*). Consider a path *q* obtained by concatenating paths Caterpillar Sort(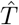, *T*) and *p*. Note that *q* is a path from 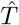 to *R* which is shorter than Caterpillar Sort(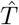, *R*). Indeed, the two paths coincide between 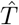 and *T*, and *q* is shorter between *T* and *R*. Since 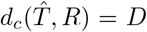, the existence of *q* is a contradiction of Lemma 4. □

### 3.2. Diameter and radius

Gavryushkin, Whidden and Matsen (2018, Theorem 7) gave an upper bound for the diameter of RNNI space, denoted by ∆(RNNI). They showed that ∆(RNNI) ≤ *n*^2^ − 3*n* − 5/8. In this section we improve that result and calculate the exact diameter and radius of the RNNI space.

#### Theorem 6. 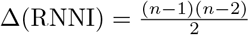

*Proof*. Since *d*_*c*_([1 …, *n*], 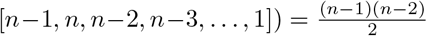, the claim follows directly from Theorem 5 and Lemma 2. □

We show in the rest of this section that the radius of RNNI space coincides with its diameter. The main tool to establish this result is the algorithm MDTree from Section 2.3. Recall that MDTree is a non-deterministic algorithm, which implies that the output tree is not uniquely defined by the input to the algorithm.

#### Lemma 7. *Let T be a caterpillar tree. Then d*(*T, R*) = ∆(RNNI) *for every tree R such that R* = MDTree(*T*)

*Proof*. We prove the lemma by induction on the number of taxa *n*. The induction basis for *n* = 3 is trivial.

Assume now that *T* = [1, …, *n* + 1]. In this case, the parent of taxon *n* + 1 has rank *n* in *T* and rank 1 in *R*. Lemma 3 then implies that

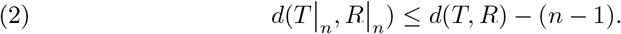

Note that *R*|_*n*_ = MDTree(*T*|_*n*_) for an appropriate choice of attachment points in the execution of MDTree. By the induction hypothesis and Theorem 6, 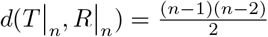. So inequality (2) becomes 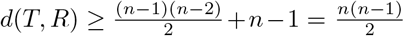, which gives the desired equality with Theorem 6. □

The non-determinism of MDTree can be exploited to investigate which trees are at the maximal possible distance from a given caterpillar tree. In fact, those trees can have an arbitrary (non-ranked) topology. This is because in each step of the algorithm the next taxon can be added on any edge incident to a leaf in the running tree. Therefore, MDTree can be used to find a tree of pre-defined topology that is at the maximal possible distance from the input tree, in particular if the input is a caterpillar tree. Because the distance between trees is invariant under permutations of taxa labels, we conclude that for every tree *R* on *n* taxa there exists a caterpillar tree *T* such that 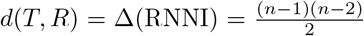. This proves the following Theorem 8.

#### Theorem 8.

*The radius and diameter of the* RNNI *space coincide and are equal to* 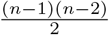

### 3.3. Cluster Property

In this section we discuss the so-called Split Property, which states that every split shared between two trees is maintained along shortest paths between the trees. For example, the SPR space has the Split Property while NNI does not (Li, Tromp and Zhang 1996). Specifically, in NNI there exist two trees *T* and *R*, which share a split *A|B*, such that every tree on every shortest path between *T* and *R* does not have *A|B*.

To construct this example, Li, Tromp and Zhang (1996) used the idea illustrated in Figure 4 and showed that for an appropriate permutation of the taxa in the two caterpillar subtrees of both *T* and *R*, it is strictly shorter to merge the two caterpillar subtrees first, then sort them simultaneously, and then split back to two caterpillar subtrees, rather the two caterpillar subtrees independently.

**Figure 4.**
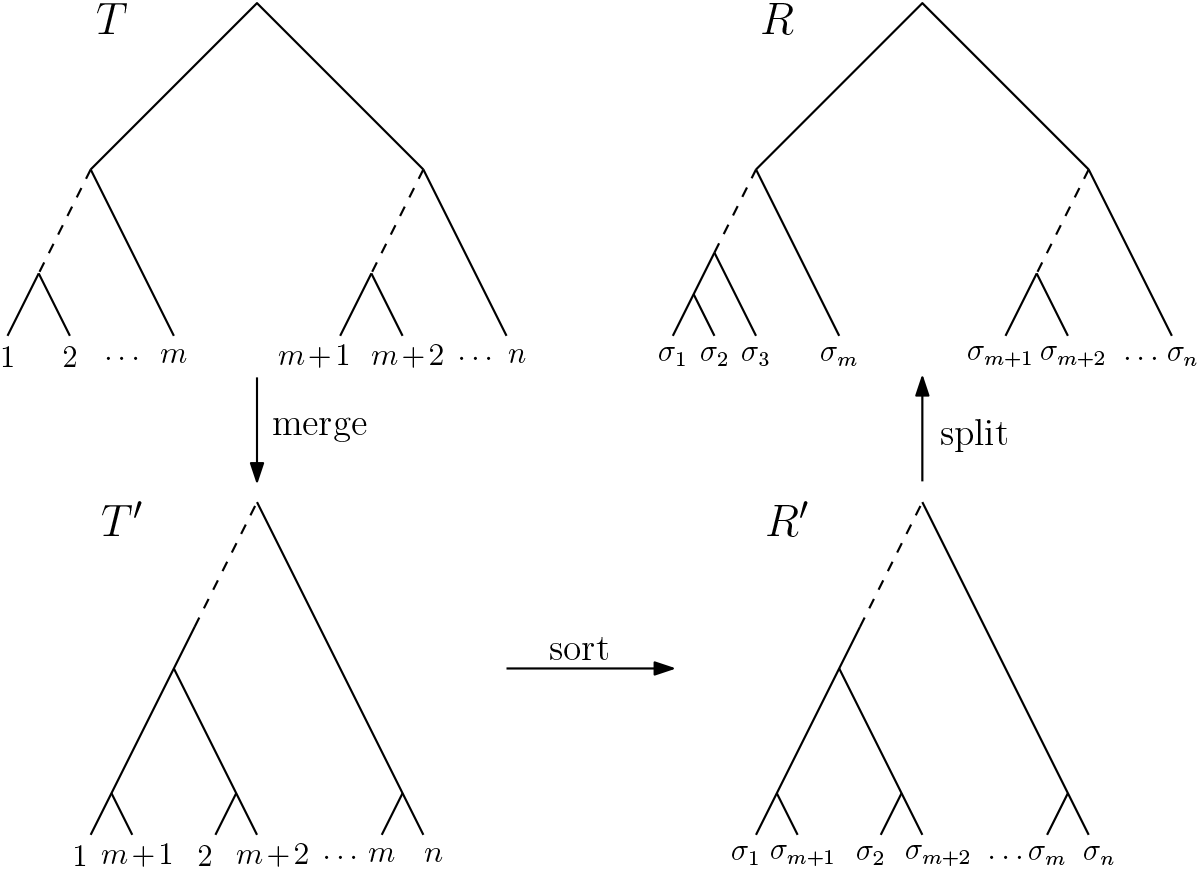
Example of a shortest path between two trees *T* and *R* in NNI. In (Li, Tromp and Zhang 1996) it is proven that there is a labelling that ensures that there is no shortest path in NNI where the clusters {1, …, *m*} and {*m* + 1, …, *n*} with 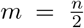 shared between *T* and *R* are preserved. Instead, the depicted path where 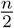 cherries are built and then sorted before being resolved is a shortest path

The reason this example cannot be transferred to RNNI is that additional rank moves are necessary to merge and then split the two caterpillar subtrees. Similarly to the example in Figure 3, adding rank moves to the shortest NNI path from *T* to *R* results in a path that is not a shortest RNNI path. Sorting the two caterpillar subtrees of *T* and *R* independently results in a shorter path in RNNI.

This argument motivated Gavryushkin, Whidden and Matsen (2018, Conjecture 9) to conjecture that every split shared between trees in RNNI is maintained along every shortest path.

The example in Figure 5 shows that, in this form, the Split Property is not present.

**Figure 5.**
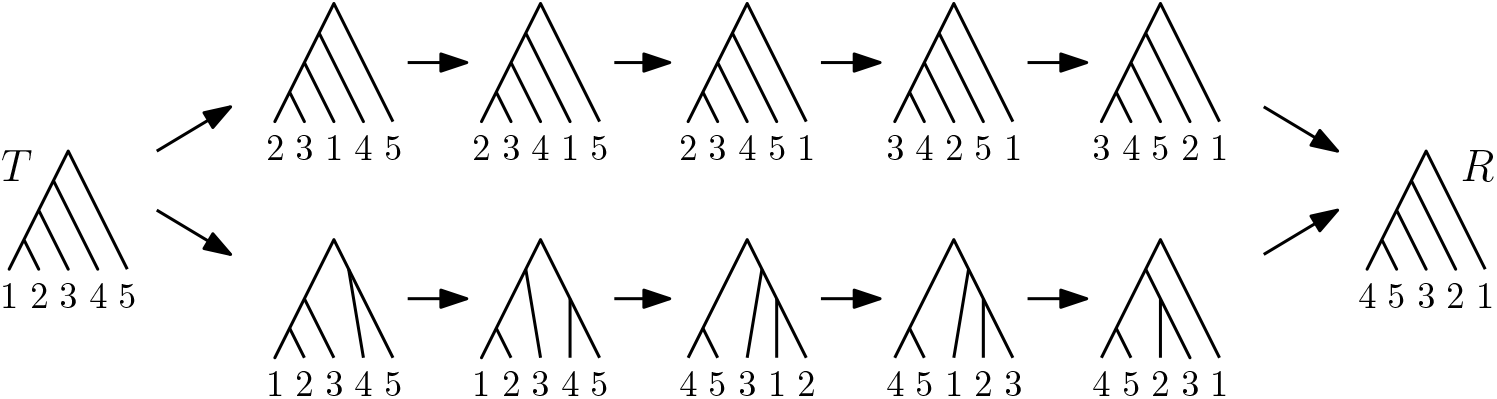
The split 123|45 is present in *T* and *R*, but the path at the top is a shortest path (computed by Caterpillar Sort) where none of the trees contains this split. On the path at the bottom, which is a shortest path as well, this split is maintained.

Hence not every shortest path between trees in RNNI has to maintain every shared split of taxa. However, there exists yet another shortest path connecting the two trees, along which every shared split is maintained. This motivates the study of the following Weak Split Property in RNNI.

#### Definition (Weak Split Property)

*If a split of taxa A|B is shared by two trees T and R then there exists a shortest path p between T and R such that all trees on p share the split A|B*.

#### Conjecture 9.

RNNI *has the Weak Split Property*.

Another interesting property of our counterexample in Figure 5 is that the set of taxa {1, 2, 3}, which is a part of the split in both *T* and *R*, does not form a cluster in *R*, that is, there is no single node in the tree all descended taxa of which are exactly 1, 2, and 3. Importantly, rooted phylogenetic trees can be uniquely represented by sets of clusters (Steel 2016), but not by sets of splits as splits cannot define the position of the root. For example, trees *T* and *R* in Figure 5 induce the same set of splits, but share no cluster.

Furthermore, all our methods in this paper for computing shortest paths in special subspaces of RNNI demonstrate that shared clusters are maintained along shortest paths.

These considerations motivate us to conjecture the following Cluster Property.

#### Definition (Cluster Property)

*Let T and R be trees that share a cluster C. Then C is present as a cluster in every tree on every shortest path between T and R.*

#### Conjecture 10.

RNNI *has the Cluster Property*.

We have found no counterexamples to this conjecture and have checked that it holds in RNNI spaces with up to seven taxa (*n* = 2, *…*, 7). Specifically, we computed subgraphs of RNNI containing only trees with shared clusters and compared distances in this subgraph with distances in the whole RNNI graph. We computed these graphs according to the algorithm from (Gavryushkin, Whidden and Matsen 2018) as discussed in Section 2. The source code of all these implementations is available at (Collienne, Elmes and Gavryushkin 2019).

### 3.4. Partition lattice

In this section we establish a connection between the RNNI graph and a well-known algebraic structure, the partition lattice. This connection provides a new direction for further research and translates results and open problems from the language of phylogenetics to the language of lattice theory.

The *partition lattice* on {1, …, *n*} is the lattice given by the partially ordered set (Π_*n*_,≤), where Π_*n*_ is the power set of {1, …, *n*} and *X ≤ Y* if partition *X* refines *Y*, that is, *X ≤ Y* ⇔ (∀*x ∈ X*)(∃*y ∈ Y*)*x ⊆ y*. For simplification we will denote the partition lattice on *n* elements by Π_*n*_. Π_4_ is illustrated in Figure 6. We assume that a partition *X* in the partition lattice Π_*n*_ has rank *k* if the number of elements in *X* is *n − k*. The algebraic structure of Π_*n*_ is related to the RNNI graph on *n*-taxa trees in the following way.

**Figure 6.**
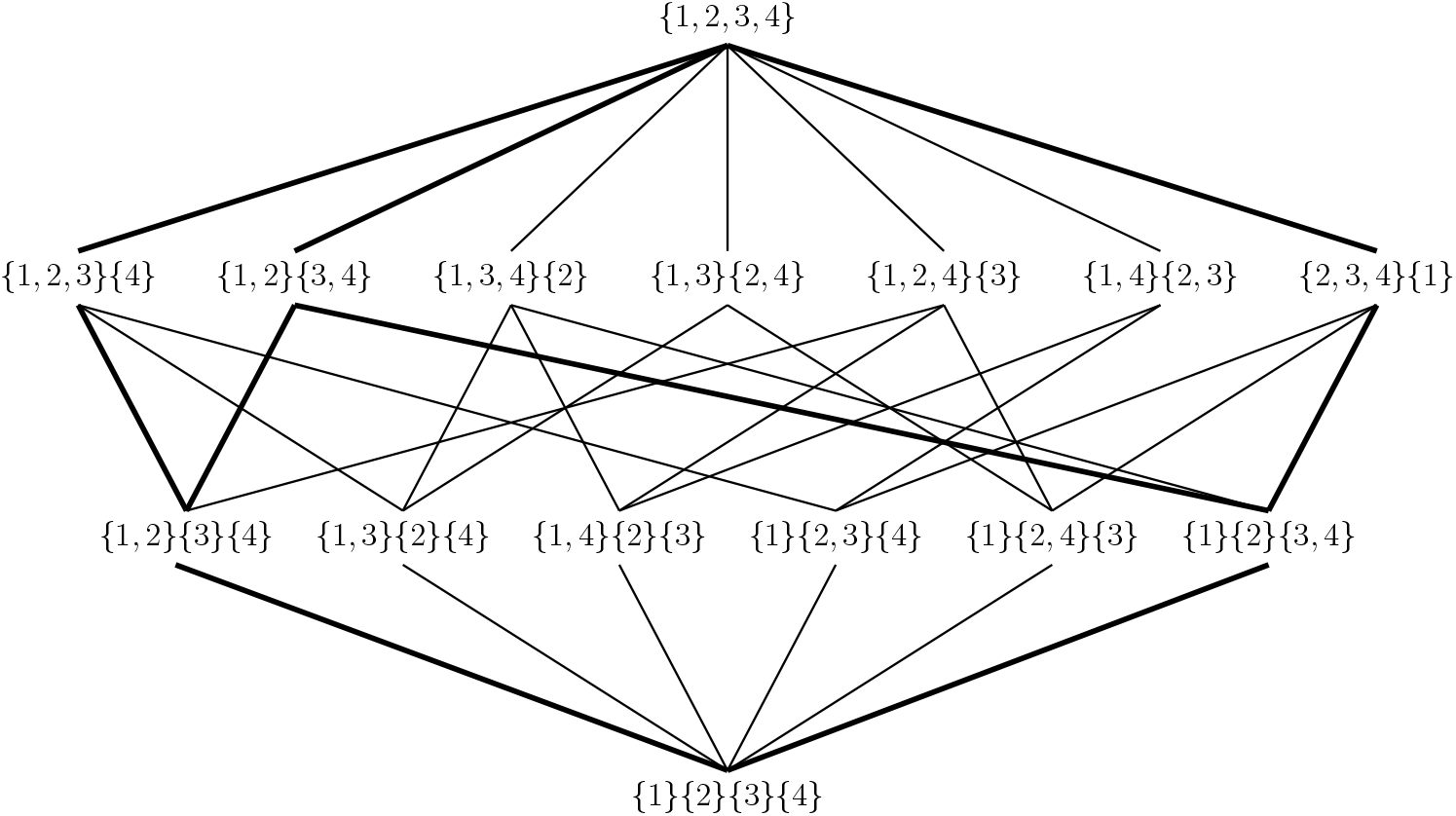
The partition lattice Π_4_ on {1, 2, 3, 4}. The highlighted edges correspond to an RNNI path from the tree represented by the leftmost chain to the rightmost one.

#### Theorem 11. *The* RNNI *graph on n taxa is isomorphic to the graph of maximal chains of the partition lattice* Π_*n*_ *where two maximal chains are connected by an edge if and only if they differ by exactly one partition. The corresponding metric spaces are isometric*

*Proof*. There is a one to one relation between ranked trees and maximum chains in a partition lattice. We can define a bijective mapping from the set of ranked trees to the set of maximum chains in Π_*n*_ as follows. A tree *T* maps onto a maximum chain *𝒞*_*T*_ if the set in the partition of rank *i* in *𝒞*_*T*_ that is the union of two sets of the partition of rank *i* − 1 in *𝒞*_*T*_ is the cluster induced by the internal node of rank *i* in *T*.

Note that this bijection is an isomorphism between the RNNI graph and the graph of chains as in the theorem. Indeed, two chains are different by exactly one partition if and only if the corresponding trees are connected by an RNNI move. □

Figure 6 is an illustration of the proof of Theorem 11. The four chains indicated in bold correspond to the following RNNI path. The leftmost chain corresponds to the caterpillar tree [1, 2, 3, 4]. First, the partition {1, 2, 3}{4} is replaced with {1, 2}{3, 4} and we get the chain corresponding to the tree [{1, 2}, {3, 4}] (in the cluster representation), which is one RNNI move away from the caterpillar tree. Second, the partition {1, 2}{3}{4} is replaced with {1}{2}{3, 4}, which corresponds to the rank swap on the previous tree. Third, the partition {1, 2}{3, 4} is replaced with {1}{2, 3, 4} and we reach the caterpillar tree [3, 4, 2, 1].

## 4. Discussion

The problem of computing distances in all common phylogenetic graphs, including NNI (Dasgupta et al. 2000), SPR (Bordewich and Semple 2005), and TBR (Allen and Steel 2001), is 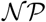-hard. Although 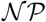-hardness does not necessarily imply practical impossibility, this so far has been the case for these phylogenetic algorithms with extremely few major advances (Whidden, Beiko and Zeh 2010). NNI is especially known for its algorithmic hardness (Whidden and Matsen 2018).

Our results suggest a surprising advance in this field of 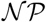-problems – adding ranking information to trees simplifies some of the algorithms and makes some unintuitive counterexamples impossible. We hence continued the investigation of the RNNI graph on ranked phylogenetic trees in this paper. We have taken the route of comparing the classic NNI with the RNNI graph, so have mostly been concentrating on questions that are settled in NNI. Specifically, we have considered a number of geometric properties, including the diameter, radius, convexity of caterpillar trees, and the Split Property. We have also justified that the technique of Dasgupta et al. (2000) to prove that distances in NNI are 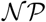-hard to compute cannot be directly applied in RNNI. This led us to the Cluster Property, a statement that does not hold in NNI. All our algorithms and methods developed in this paper witness in favour of RNNI having the Cluster Property. For example, we have checked the statement computationally for up to seven taxa (Collienne, Elmes and Gavryushkin 2019). Although the Split Property is present in SPR, it is still 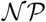-hard to compute distances in that graph. Hence the Cluster Property alone cannot be used as an argument in favour of the distance problem having an efficient solution in RNNI. But the greedy nature of all our algorithms for computing shortest paths in RNNI developed in this paper does provide such an argument.

The algorithms we developed and implemented in this paper have served as a main ingredient for our study of the geometry of RNNI space. These algorithms are of interest on their own as well. For example, our FindPath algorithm gives a good approximation of the RNNI distance, performing exactly on small trees. We are not aware of the minimal number of taxa where this algorithm fails to return the correct distance. The MDTree algorithm produces trees as far away from a given tree as possible, a task of importance in simulation and model comparison studies. The Caterpillar Sort algorithm efficiently computes the RNNI distance and shortest paths between caterpillar trees exactly. To the best of our knowledge no such algorithm exists for NNI. This implies that the algorithmic complexity of computing distances in NNI is higher than in RNNI.

The question of whether computing RNNI distances is 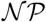-hard is still open.

## Acknowledgements

We thank Charles Semple for useful discussions about the Cluster Property, and Mike Steel for his useful comments that improved this paper.

We acknowledge support from the Royal Society of New Zealand through the Rutherford Discovery Fellowship (RDF-UOO1702) awarded to AG. This work was partially supported by the Ministry of Business, Innovation, and Employment of New Zealand through the Endeavour Smart Ideas (CONT-61378-ENDSI-UOO) and Data Science Programmes grants.

MF thanks the joint research project *DIG-IT!* supported by the European Social Fund (ESF), reference: ESF/14-BM-A55-0017/19, and the Ministry of Education, Science, and Culture of Mecklenburg-Vorpommern, Germany, for funding parts of this work.

Part of this work was done while AG, LC, and MF were visiting Max Plank Institute for Evolutionary Biology, we appreciate their support.

